# Towards real-world neuroscience using mobile EEG and augmented reality

**DOI:** 10.1101/2021.09.29.462177

**Authors:** Alexandra Krugliak, Alex Clarke

## Abstract

Our visual environment impacts multiple aspects of cognition including perception, attention and memory, yet most studies traditionally remove or control the external environment. As a result, we have a limited understanding of neurocognitive processes beyond the controlled lab environment. Here, we aim to study neural processes in real-world environments, while also maintaining a degree of control over perception. To achieve this, we combined mobile EEG (mEEG) and augmented reality (AR), which allows us to place virtual objects into the real world. We validated this AR and mEEG approach using a well-characterised cognitive response - the face inversion effect. Participants viewed upright and inverted faces in three EEG tasks (1) a lab-based computer task, (2) walking through an indoor environment while seeing face photographs, and (3) walking through an indoor environment while seeing virtual faces. We find greater low frequency EEG activity for inverted compared to upright faces in all experimental tasks, demonstrating that cognitively relevant signals can be extracted from mEEG and AR paradigms. This was established in both an epoch-based analysis aligned to face events, and a GLM-based approach that incorporates continuous EEG signals and face perception states. Together, this research helps pave the way to exploring neurocognitive processes in real-world environments while maintaining experimental control using AR.

## Introduction

A great deal of progress has been made in cognitive neuroscience, where imaging techniques offer a window into the mysteries of visual perception, memory, attention and language, to name a few. Such success has largely been achieved with a scientific approach where researchers seek to isolate specific cognitive functions and study their neurocognitive instantiation in a controlled manner. This is an important and fruitful approach, and will continue to be so. As cognitive neuroscience methods progress, a complementary approach has also become more feasible, namely, research using more naturalistic paradigms which look towards the functioning of the human mind unleashed from controlled experiments (Aliko et al., 2020; Allen et al., 2021; Jääskeläinen et al., 2021; Matusz et al., 2019). Moreover, experimentation beyond the lab is possible by building on the platform established by prior approaches, and by employing newer and emerging technologies to study the human brain - such as virtual reality, augmented reality and fully mobile approaches for neural recordings. Going further, combining these approaches could offer a toolkit that allows for the study of neural function in uncontrolled complex visual environments, getting us closer to studying cognition in our natural habitat. Here, we present and validate an approach to studying human cognition in naturalistic, real-world environments, while importantly retaining the ability to manipulate our key variables and retain experimental control. We achieve this by combining mobile EEG (mEEG) with head-mounted cameras and augmented reality (AR).

Mobile whole-head EEG applications have made great strides in the past decade, recently being applied to study memory (Griffiths et al., 2016; Park & Donaldson, 2019; Piñeyro Salvidegoitia et al., 2019), emotion (Packheiser et al., 2021; Soto et al., 2018), attention (Ladouce et al., 2019; Liebherr et al., 2021) and movement (Mustile et al., 2021; Packheiser et al., 2020; Reiser et al., 2021) in complex real-world settings. Advances in hardware and software have made fully mobile high-density EEG a useful tool for cognitive neuroscience (Klug & Gramann, 2021; Symeonidou et al., 2018), however, what remains problematic is the ability to flexibly manipulate key variables, as now our cognitive variables of interest are part of the real world. For some disciplines, this could be circumvented by placing certain objects in certain places, thereby constructing experimentally useful environments (e.g. Park & Donaldson, 2019). However, a more adaptable and flexible approach is to utilise immersive head-mounted displays to present virtual objects on the background of the real world, allowing for full experimental control over what people see and where those items are located. In contrast to virtual reality, where the whole environment is simulated, AR affords the ability to place 3D virtual objects in the actual environment. Recent research suggests that AR is more engaging than VR, with an indication of improved memory performance (Maidenbaum et al., 2019). The ability to place a limitless set of virtual items in the real world offers a degree of experimental control that can’t be matched through brute force methods. Here, we propose that by combining mobile EEG with head-mounted AR, we can study human cognition and neuroscience in real-world settings while also manipulating the world people see.

In order to demonstrate such an approach, we chose a well characterised and robust neural signature, the face inversion effect (Rossion et al., 1999). Participants completed three face inversion sessions while we recorded 64-channel EEG from a mobile system. First a computer-based face inversion task, second viewing photographs of faces attached to the walls of a corridor, and third viewing virtual faces through the head mounted AR device. In both the second and third task, participants were freely navigating in an indoor corridor setting. As a follow on, we next established routines whereby we could relate the dynamically unfolding visual environment to the dynamic neural signatures recorded while participants walked through natural environments. This is important, as to be able to study cognition in real-world settings, we need methods to link natural dynamic behaviour to dynamic neural signals. To achieve this, we used a GLM approach similar in nature to that used in naturalistic fMRI studies of movie watching (e.g. Hasson et al., 2004; Huth et al., 2012) and MEG studies of language comprehension (e.g. Brodbeck et al., 2018), again testing the sensitivity of the approach against face inversion effects. Together, these twin analyses demonstrate an approach that manipulates and controls variables in conjunction with mobile neural recordings to reveal cognitive effects in dynamically changing settings.

## Methods

### Participants

Eight healthy participants (age 19-38 years, 4 females) took part in the study, and all had normal or corrected to normal vision. The experiment was approved by the Cambridge Psychology Research Ethics Committee and all participants gave informed consent.

### Stimuli & Procedure

The experiment had three distinct tasks.

#### Task 1, Computer-based

A computer-based face processing task was conducted using upright and inverted images of a male and female face (Figure 1A). The two face images were obtained from the Psychological Image Collection at Stirling (pics.stir.ac.uk). Face images were presented in colour, were front facing on a white background and had a neutral expression. Each face was shown in both upright and inverted orientations. Participants were instructed to press a button on the keyboard when they had seen the face. Each trial began with a fixation cross lasting between 500 and 525 ms, followed by the face image lasting until the button press, before a blank screen lasting 1 second. A total of 200 trials were presented, 100 upright and 100 inverted faces. The experiment was presented using Psychtoolbox version 3 and Matlab R2019b, and triggers recorded via a USB to TTL module (https://www.blackboxtoolkit.com).

**Figure 1.**
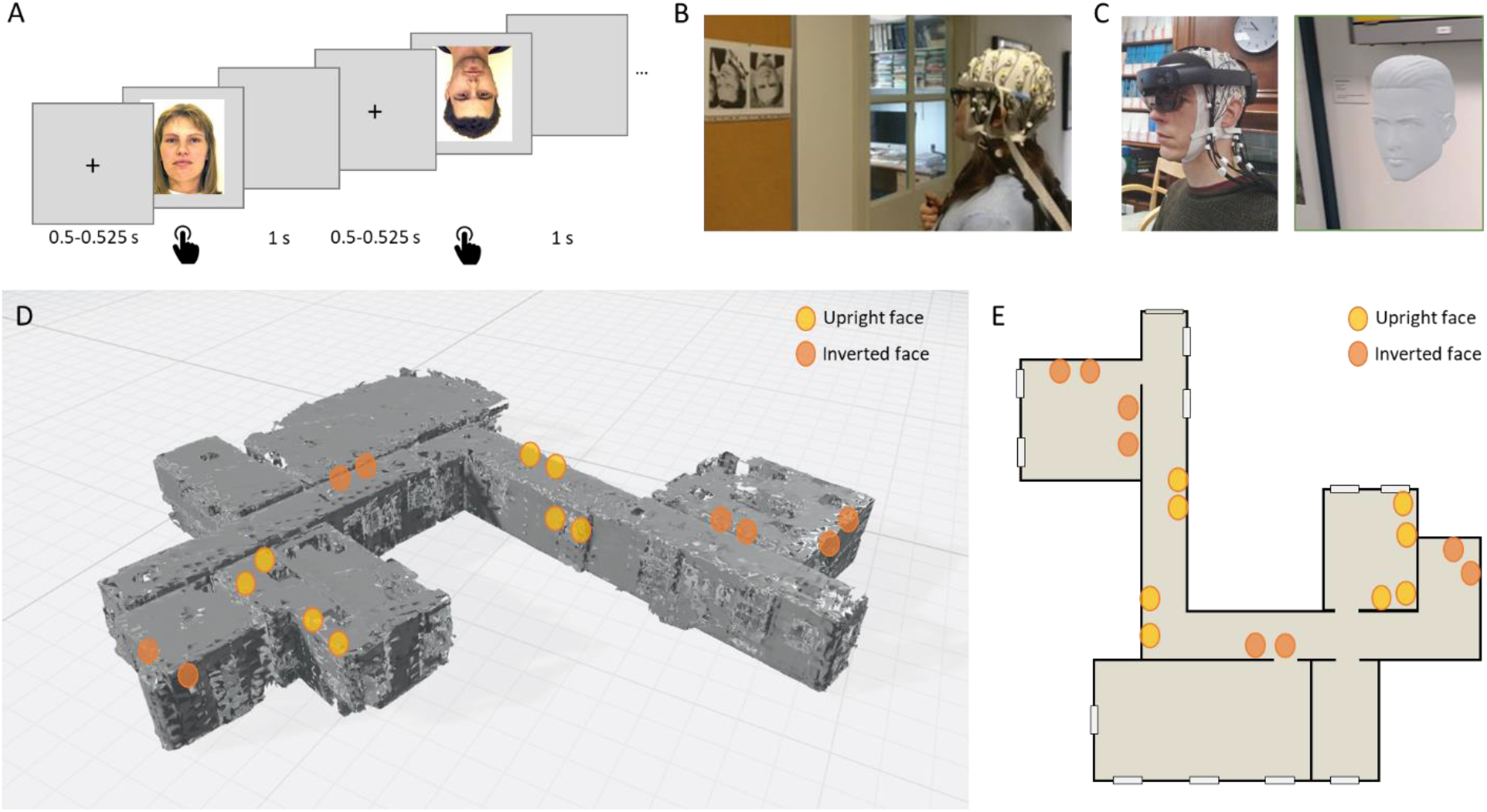
Experimental setup. A) Computer-based task showing example trials and timings. B) Mobile EEG setup and face photos. C) Mobile EEG and AR setup and example virtual face used for AR task. The photographs in B and C show the authors, used with permission. D) 3D spatial map created by the Hololens 2 of the experimental environment, showing approximate locations of upright (yellow) and inverted (orange) virtual faces. E) 2D map of environment showing approximate locations of upright (yellow) and inverted (orange) virtual faces.

#### Task 2, mEEG + photos

For task 2, photos of upright or inverted faces were attached to the walls of the corridor and participants were asked to view the faces while walking along the corridor (Figure 1B). Sixteen black and white photographs of faces (8 upright, 8 inverted) were taken from the set used by Greene & Hodges (1996). Faces were forward facing with natural expressions and plain backgrounds. In addition to EEG, participants were fitted with a head-mounted camera attached to a Raspberry Pi Zero (https://www.raspberrypi.org/). Participants were asked to move along the corridor, and into side rooms, and view each face image. When they were viewing the image they pushed a button which sent a signal to the LiveAmp trigger input. The condition the image belonged to could then be derived from the head-mounted video. Participants repeatedly viewed the face images resulting in an average of 42 upright (range 24-70) and 39 inverted trials (range 22-72). The variable number of trials across participants reflects the different routes participants chose to take.

#### Task 3, mEEG + AR

In task 3, a Microsoft Hololens 2 AR device (https://www.microsoft.com/en-us/hololens) was positioned over the electrodes (Figure 1C). The clear lenses of the Hololens allow the participant to see the actual environment, while virtual faces were presented anchored to specific locations in the corridor (Figure 1DE). The onboard camera of the Hololens captured first-person video, including the virtual faces. The virtual faces were obtained from the Microsoft 3D objects library, and were white heads with texture and hair, but no colour (Figure 1C). In this respect, they resembled white marble heads without a body. Four different virtual heads were used (2 male, 2 female), with each head appearing in 4 locations, half of which the faces were inverted. This resulted in a total of 16 heads, 8 of which were upright and 8 inverted. The virtual heads were placed along the same corridor and side rooms as used for the face photographs (which were not present during task 3). Participants pushed a button when viewing the virtual face, and saw each face numerous times resulting in an average of 63 upright (range 28-90) and 53 inverted trials (range 27-87).

### EEG recording

EEG was recorded using the Brainvision LiveAmp 64 mobile system (Brain Products GmbH, Gilching, Germany). In all tasks we recorded 64-channel EEG through ActiCap Slim active Ag/AgCl electrodes referenced to an electrode placed at FCz, with a sampling rate of 500 Hz. Electrodes were embedded in an elastic cap with electrode locations conforming to the international 10/20 system. Electrode cables were carefully routed through the cable ties and kept flat to the head to minimise cable sway during recording. EEG signals were amplified using two wireless amplifiers and recorded to onboard memory cards. Data recording was controlled using BrainVision Analyzer, and during setup, impedances were reduced to below approximately 5-10 kOhm.

### Epoch-based analysis

The same EEG analysis pipeline was independently used for the data from all three tasks, with a slight modification for the mobile tasks (2 and 3; Figure 2). Raw EEG signals were imported into EEGlab (Delorme & Makeig, 2004) and cropped to just before the first trial and just after the last trial, before channel locations and names were imported. The data were then band-pass filtered using a onepass zerophase Blackman-windowed sinc FIR filter with transition width 0.1 Hz and order 27500. Band-pass were 0.5 to 40 Hz for task 1, and 1 to 20 Hz for task 2 and 3. The narrower filter range for task 2 and 3 was due to the additional noise suppression this afforded, and the frequency range of interest identified in task 1. Bad channels were detected based on automated procedures (pop_rejchan.m), and a channel was classified as bad based on probability (threshold 3 SDs), spectrum (threshold 3 SDs) and kurtosis measures (threshold 5 SDs). Any identified bad channels were subsequently interpolated using spherical interpolation (task 1: mean=9.4, range=7-14; task 2: mean=5.9, range=1-11; task 3: mean=7.8, range=4-12). The continuous data was then cleaned using clean_artifacts. For task 1, the data were epoched between −1 and 2 seconds around the onset of the face images, and for task 2-3 the data were epoched between −2 seconds and +2 seconds centred on the button press. The condition of each epoch was then determined using the head-mounted videos. The epoched data was visually inspected and noisy trials were removed (task 1: mean=10%, range=0-47%; task 2: mean=7%, range=0-14%; task 3: mean=9%, range=1-19%).

**Figure 2.**
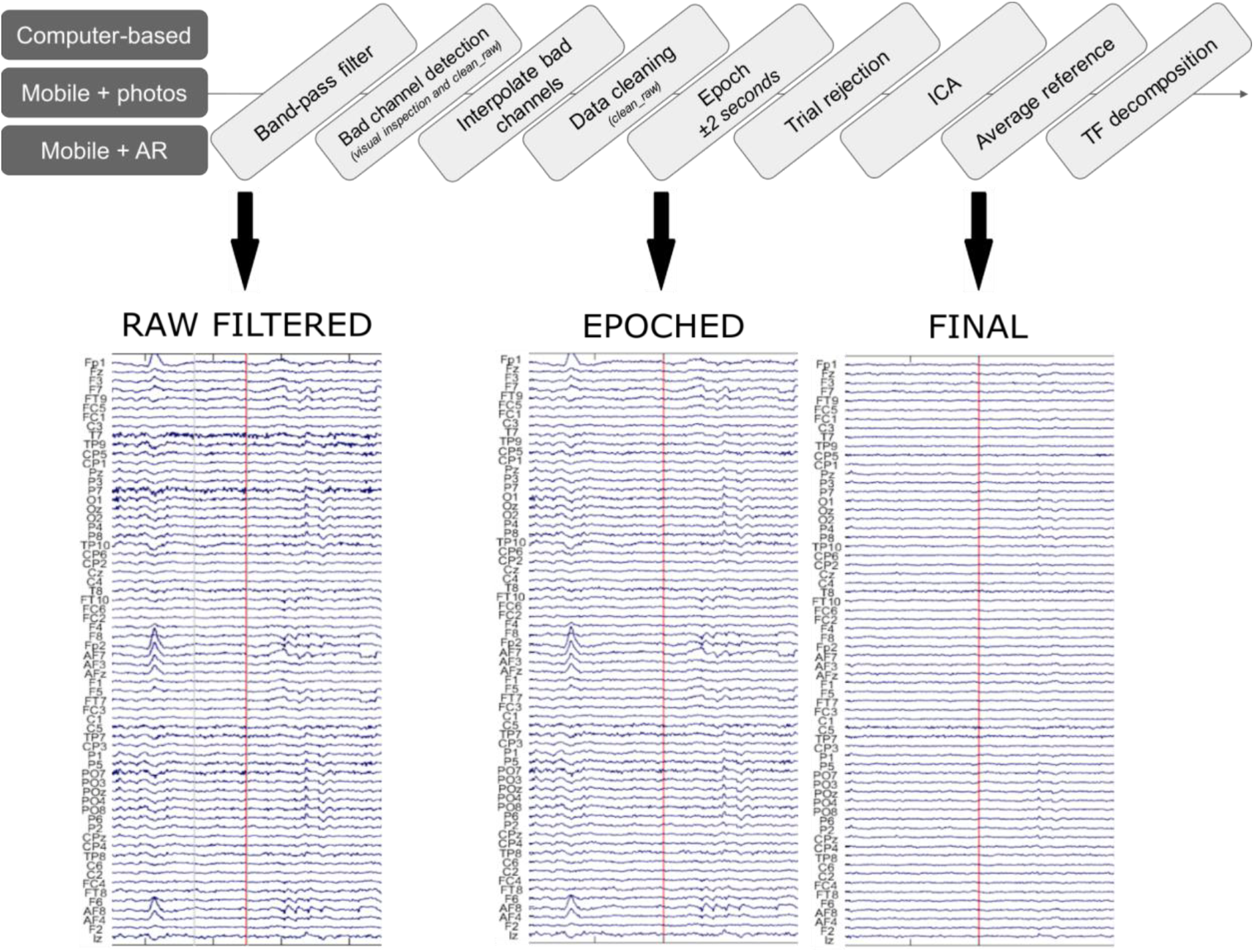
*EEG preprocessing. Top, core methods across* task, *and example data at different stages of processing from the Mobile + AR task*.

We used ICA to focus our analysis on signals coming from the brain rather than external or noise sources. ICA was applied to the epoched data using runica with the extended and pca options, and extracted N components where N was 64 minus the number of bad channels. ICs were analysed using ICLabel (Pion-Tonachini et al., 2019) to identify components related to brain activity to retain when reconstructing the EEG data. Components were retained if they had a greater than 20% chance of reflecting brain activity but could not have more than 20% chance of reflecting any artefact classification (eyes, cardio, muscle, channel noise). These components were then visually inspected and any remaining components not showing a 1/f characteristic or with only frontal electrode weights were additionally removed. On average 13.9 components were retained in task 1 (range 5-30), 6.4 in task 2 (range 2-11) and 8.3 in task 3 (range 4-13). We used this very selective approach to help focus the EEG signals on cognitively relevant signals. After ICA, the data were again visually inspected before transforming the signals using an average reference. The processed EEG signals were converted to Fieldtrip (Oostenveld et al., 2010), and time-frequency representations of each trial were calculated using Morlet wavelets between 4 and 35 Hz in 25 ms time steps between −0.25 and 1.25 seconds (task 1), or 4 to 20 Hz in 25 ms time steps between −1.5 and 1.5 seconds (task 2 and 3). No baseline correction was applied.

We contrasted low frequency oscillatory power averaged between 5 and 15 Hz based on prior studies showing enhanced power for inverted compared to upright faces in this range. Further, as these effects are expected over posterior electrodes, we initially focused our analysis on these electrodes. To do this, power was averaged across posterior electrodes (N=17, defined as all occipital, parietal and parietal-occipital electrodes) and between 0 to 400 ms (task 1) and −1.5 to 1.5 seconds (task 2 and 3). This produced one value per trial before the data were converted to a standardized z-score. A linear mixed effects model was used to model trial-wise power with a predictor variable specifying if the trial was an upright face (−0.5) or an inverted face (0.5), and including a random effect of Subject. Trials with a value more than 4 standard deviations away from the mean were additionally removed prior to the mixed effects model (task 1: 0.5% of data; task 2: 0.2%; task 3: 0.2%).

### GLM-based analysis

The GLM-based approach was applied to the mobile tasks (2 and 3). We took the ICs selected for the epoch-based analysis, and applied them to the filtered data from each task. We then created an additional stimulus channel in the data, by inserting a stick function at the time of each face event with values rising to 0.5 for an inverted face and −0.5 for an upright face. All other time points were set to zero. This stimulus channel was then convolved with a 2 second hamming window centred on the face events. The continuous EEG was then visually inspected, and noisy time segments were selected and cropped from the data (task 2: mean=1.3%, range 0.1-4.7%; task 3: mean=7.4%, range=0-17.3%).

The EEG was imported into Fieldtrip (Oostenveld et al., 2010), and a time-frequency representation of the data were calculated using Morlet wavelets between 4 and 20 Hz in 25 ms time steps. No baseline correction was applied, and time-frequency signals were averaged across 5 to 15 Hz (following the epoch-based analysis). This resulted in a low frequency power time-series at each electrode. To test for face inversion effects over posterior electrodes, a general linear model was run fitting the power averaged over posterior electrodes with predictors of the face inversion channel, and three additional predictors of no-interest defined by the three accelerometer channels (amplifier motion in x, y and z directions) using fitglm in Matlab. Prior to the GLM, data were converted to a standardized z-score.

### Participant motion and EEG

Finally, we quantified how participant motion, measured by the accelerometers within the amplifiers which were attached to the participants back, related to EEG signal amplitudes. To do this, the continuous data (EEG channels plus 3 accelerometer channels) were split into 2 second non-overlapping chunks, and the root mean square (RMS) calculated for each channel. The RMS was averaged across the three accelerometer channels, creating one value per 2 second period. These values were then binned into low, medium and high motion RMS groups using the discretize function in Matlab, before averaging the EEG RMS according to the motion RMS bins. This resulted in an EEG RMS value for each electrode and each of the low, medium and high RMS bins. A linear mixed effects model was used to test if the EEG RMS values related to the levels of participant motion (as defined by the motion RMS bins).

## Results

### Epoch-based analysis

#### Task 1: Computer face inversion task

Our initial analyses looked to replicate the well-characterised face inversion effect in a standard laboratory setting. We calculated low frequency power for upright and inverted face trials for the first 400 ms of the face appearing, averaged the power across 5 to 15 Hz, and averaged these values across a set of posterior electrodes. To test for changes in power for upright and inverted faces, linear mixed effects modelling was used, showing that EEG low frequency power over posterior electrodes was significantly greater for inverted faces compared to upright faces (mean difference = 0.404, t(1427) = 8.27, p < 0.0001; Figure 3A). This result is expected, and replicates previous reports (Olivares et al., 2015; Tang et al., 2008). Plotting EEG power across frequencies further indicated that these increases in low frequency power peaked near 10 Hz, and differences between inverted and upright faces were principally between 5 and 12 Hz (Figure 3B). Finally, to examine face inversion effects beyond the posterior electrodes, inversion effects were calculated for each electrode, showing greater power for inverted compared to upright faces that were primarily located at posterior central and posterior lateral electrodes (Figure 3C).

**Figure 3.**
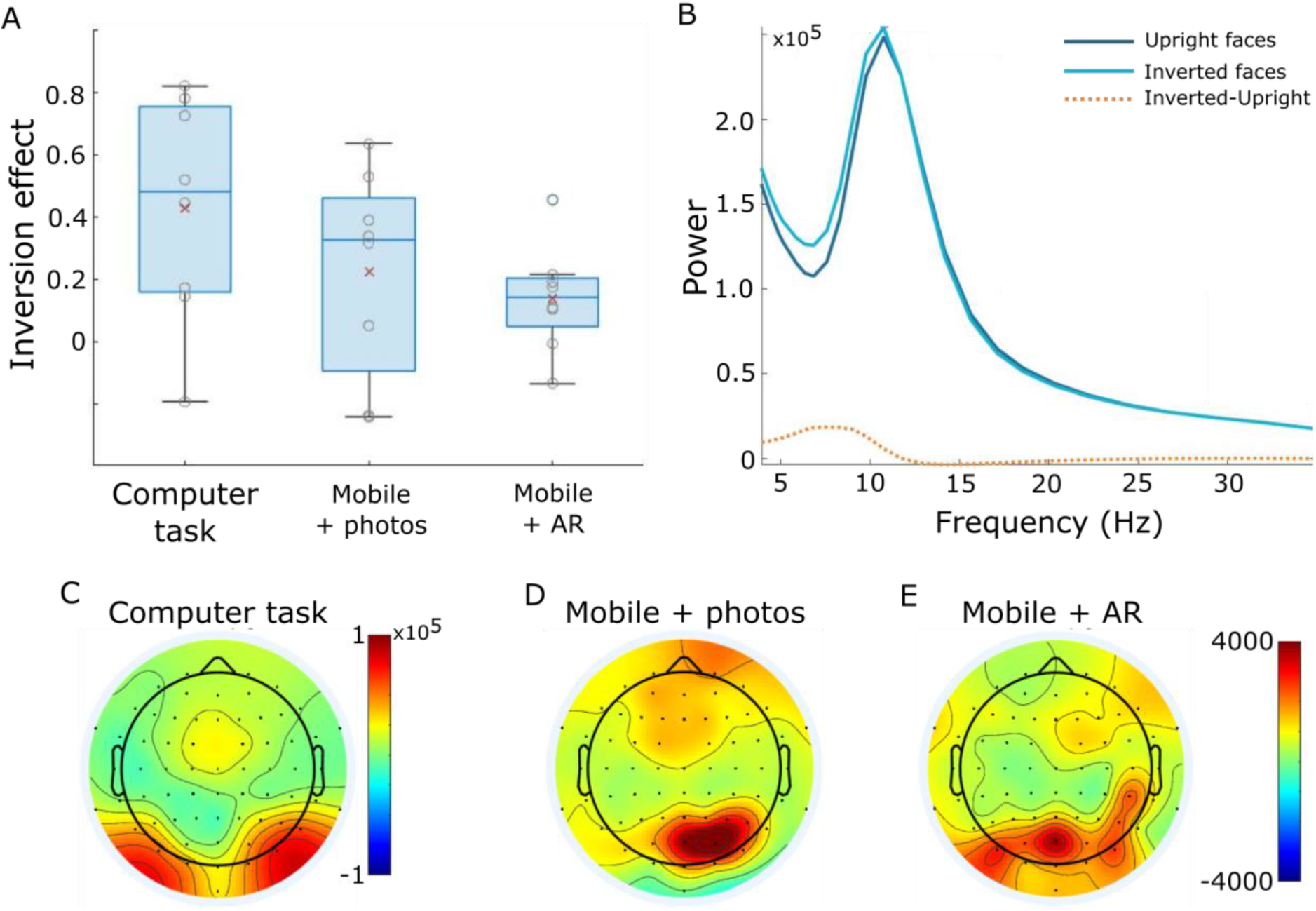
Epoch-based results. A) Face inversion effect sizes for each experimental task based on the posterior electrodes. Red x indicates group mean inversion effect with individual subjects shown by grey circles. B) Spectrogram showing group mean power between 4 and 35 Hz for upright and inverted conditions, and the difference between them. C-E) Topographies showing mean power difference for inverted-upright faces between 5 and 15 Hz for the computer task (C), mEEG + photos (D) and mEEG + AR (E).

#### Task 2: mEEG + photos

Our next analysis asked whether face inversion effects could be seen in a more naturalistic setting using mobile EEG while participants viewed upright and inverted pictures of faces placed on the walls. Black and white photos of faces were placed on the wall at various locations along a corridor and in the adjoining rooms. Participants were fitted with the mobile EEG system and a head-mounted camera, and repeatedly walked along the corridor viewing the faces, pushing a button when they were looking at each face image. We used these button presses to segment the continuous EEG recording into 4 second epochs centered on the button press, and extracted the mean low frequency power for each face event.

Following the analysis used for the computer-based task, we first performed a linear mixed effects analysis EEG power averaged across posterior electrodes. This revealed a significant effect of face inversion, with greater power for inverted faces compared to upright faces (mean difference = 0.190, t(597) = 2.38, p = 0.017; Figure 3A). Across the scalp, inversion effects were greatest over posterior central electrodes and frontal electrodes (Figure 3D). Effect sizes were maximum over posterior electrodes, and were approximately half the size of the effect sizes seen during the computer-based task (Figure 3A). This shows that face inversion effects are detectable using mobile EEG in a natural indoor setting.

#### Task 3: mEEG + AR

An important issue for studies using mobile EEG in natural settings, is the ability to manipulate the environment for the purposes of the experiment. Here, we combine mobile EEG with a head-mounted AR system which enables us to present virtual objects embedded within the real environment. Using the same corridor setting as for task 2, upright and inverted virtual heads were placed at various locations along the corridor and in the adjoining rooms. Like in task 2, participants repeatedly viewed the faces and pressed a button when they were fixating on the face. Again, the continuous data were segmented using the timings from the button presses, creating 4 second epochs centred on the button press, before calculating low frequency power for each face event.

A linear mixed effects model of EEG power averaged over posterior electrodes showed a significant effect of face inversion, with greater power for inverted compared to upright faces (mean difference = 0.170, t(828) = 2.54, p = 0.0112; Figure 3A). Across the scalp, the inversion effect was maximal over posterior central electrodes (Figure 3E). These effects were partially overlapping with those seen in the computer-based and Mobile+photos tasks, with similar effect sizes in both mobile tasks. Through the combination of mobile EEG and head-mounted AR, this analysis establishes a feasible approach to studying cognitive processes in natural, real environments in which the participant is immersed.

### GLM-based analysis

While the epoch-based analyses establish that mobile EEG and AR are a feasible combination for cognitive neuroscience, an important step is being about to relate the changing visual world to dynamic neural signals in a more flexible manner. This can be achieved through relating the time-varying EEG signals to a time-varying signature reflecting cognitive events. As such, we employed an approach to link face inversion to the EEG signals reminiscent of research relating BOLD responses to visual events using a GLM approach (Hasson et al., 2004; Huth et al., 2012).

GLMs including a face inversion regressor were run for each participant (Figure 4). For the Mobile + photos task, EEG low frequency power was averaged across posterior sensors prior to the GLM. This revealed a significant effect of face inversion, with greater power for inverted faces compared to upright faces (mean difference = 0.128, t(7) = 3.71, p = 0.0076; Figure 5A). In addition, effect sizes were calculated at each electrode which showed greater power for inverted compared to upright faces over posterior electrodes (Figure 5B). For the Mobile + AR task, the GLM based on posterior sensors also revealed a significant effect of face inversion, with greater power for inverted faces compared to upright faces (mean difference = 0.126, t(7) = 2.7, p = 0.031; Figure 5A). Additionally, face inversion effects were greatest over posterior electrodes (Figure 5C). It is also notable that the effect size maps across electrodes are highly similar for the epoch-based and GLM-based analyses.

**Figure 4.**
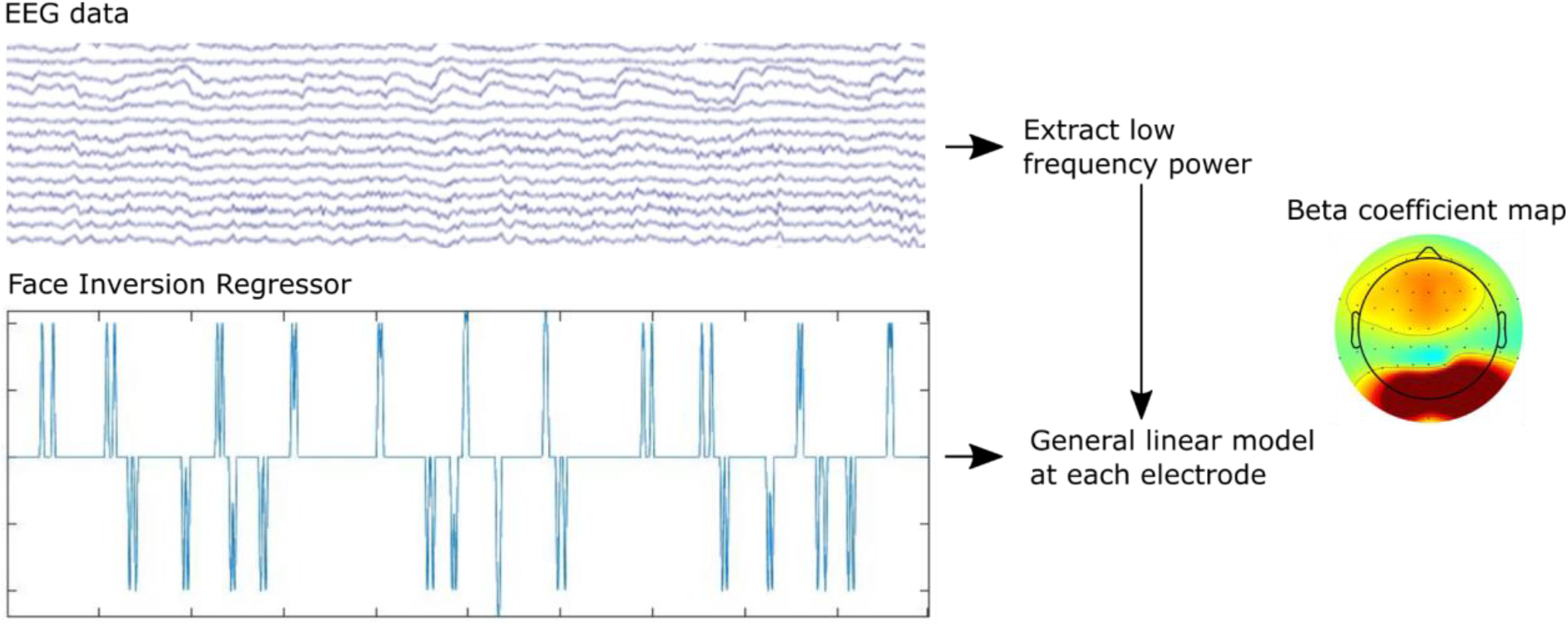
GLM-based approach. The preprocessed, continuous EEG signals were converted to low frequency power over time. A face inversion channel, the same length as the EEG data, was created where a positive spike occurred when an inverted face was seen, and a negative spike occurred when an upright face was seen. This channel was convolved with a 2 second hamming window to create the face inversion regressor. The low frequency EEG signals were modelled by the face regressor using a general linear model at each electrode, resulting in a beta coefficient map for each participant.

**Figure 5.**
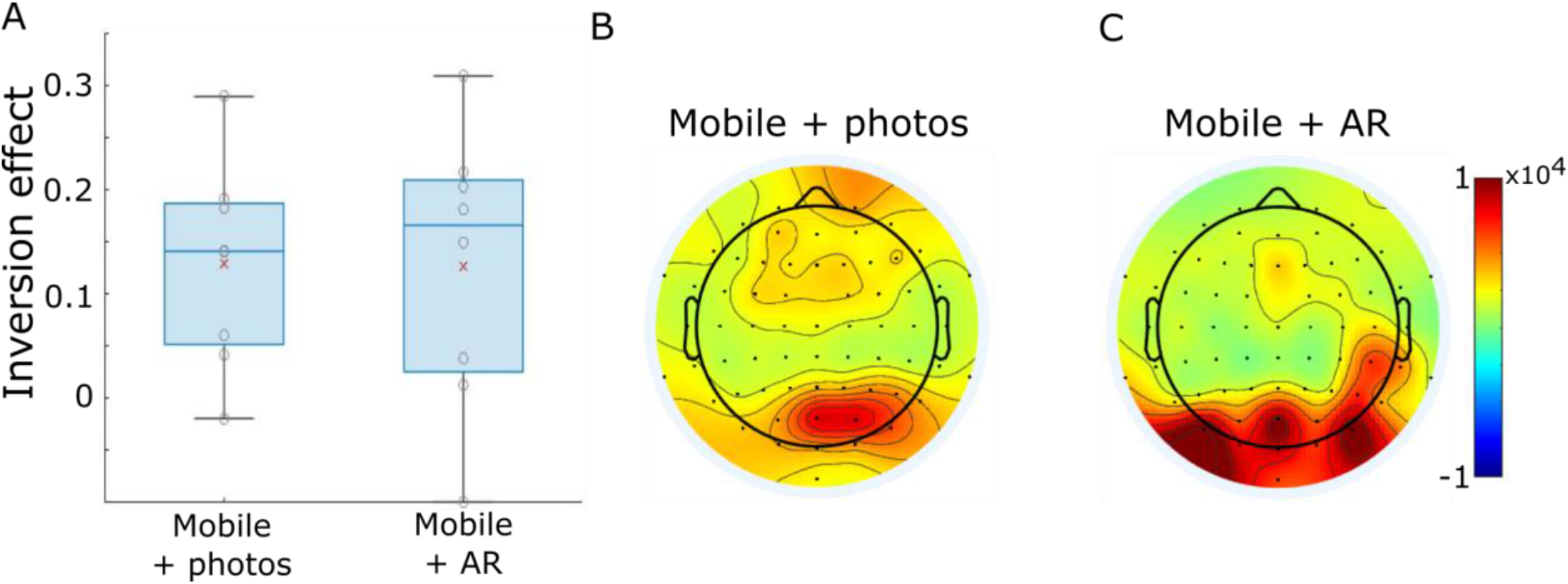
GLM-based results. A) Face inversion effect sizes for the mobile experimental tasks based on the posterior electrodes. Red x indicates group mean inversion effect with individual subjects shown by grey circles. B) Topography showing mean power difference for inverted-upright faces between 5 and 15 Hz for the mEEG + photos task, and (C) the mEEG + AR task.

In summary, we show that combining continuous event markers of perception, with continuous mobile EEG can reveal face inversion effects similar to those obtained using an epoch-based approach and contrasting conditions. This provides an approach that can be very flexible whereby multiple events and event types could be simultaneously modelled during mobile EEG.

### Participant motion and EEG

As a final assessment of the EEG signals during mobile recording, we assessed whether information from the EEG systems accelerometer channels related to signals from the EEG electrodes. To do this, we split the continuous data into 2 second non-overlapping segments, and calculated the root mean square (RMS) for the accelerometers (motion RMS) and each EEG electrode. Motion RMS was divided into low, medium and high groups, and related to EEG RMS using a linear mixed effects model. EEG RMS signals appear stable over the 3 motion levels for both mobile tasks (Figure 6AB), with no significant effects of motion level on EEG RMS (Mobile + photos: estimate −0.003, t(1534) = −0.12, p = 0.90; Mobile + AR: estimate 0.022, t(1534) = 0.79, p = 0.43). Note, equivalent results are present using a measure of signal variance rather than RMS.

**Figure 6.**
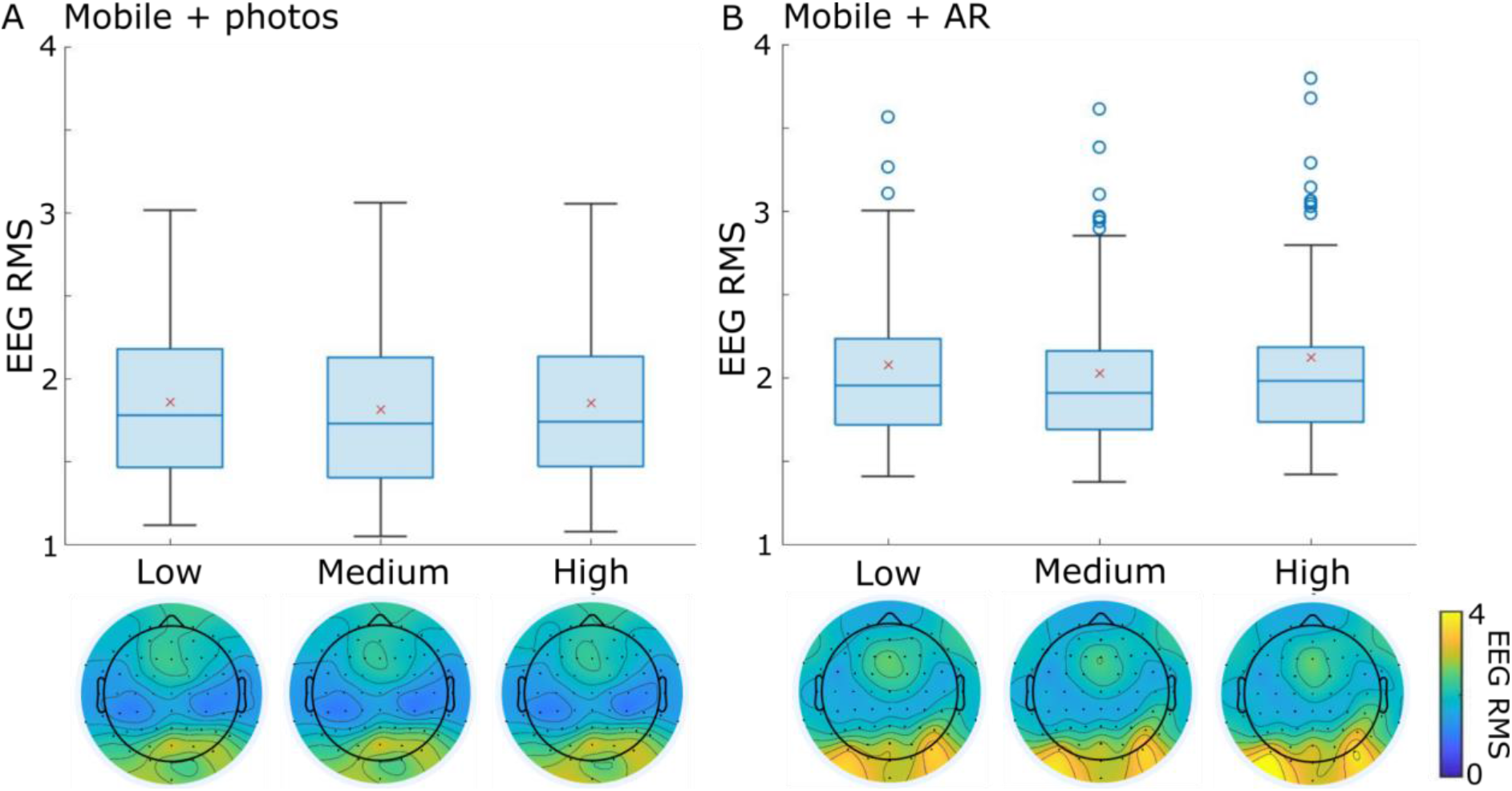
EEG RMS during low, medium and high participant motion for A) the Mobile + photos task and B) the Mobile + AR task. Red x indicates mean RMS over electrodes and participants with boxplots showing the distribution of RMS values across electrodes. Blue circles show outlier electrodes based on RMS values (not motion). Topographies show electrode RMS values for each motion RMS bin.

## Discussion

In the current study, we combined mobile EEG and head-mounted AR to establish a feasible approach to studying cognitive processes in natural, real environments in which the participant is immersed. In order to establish the feasibility of our approach, participants completed three EEG face inversion tasks: (1) a computer-based task, (2) a mobile task with photographs of faces on the walls, and (3) a mobile task where virtual faces were presented through the head-mounted AR device. In all tasks, and in both an epoch-based analysis and a GLM-based analysis that uses continuous EEG data, we see increased low-frequency power over posterior electrodes for inverted faces compared to upright faces, replicating known face inversion effects (Olivares et al., 2015; Tang et al., 2008) but in a novel experimental paradigm. Importantly, our analyses clearly show face inversion effects can be identified during free moving EEG paradigms. Our research shows that combining whole-head mobile EEG and head-mounted AR is a feasible approach to studying cognitive processes in natural and dynamic environments, which could help open the door to studying a variety of cognitive factors in real environments, whilst also allowing for the control of visual aspects of those environments using AR.

In order to successfully analyse perception during mobile EEG, we departed from typical EEG analyses. In most EEG paradigms there is an event with a definite onset, for example when a picture might appear on screen having previously not been present. This would be akin to most lab-based studies of perception, and our data from task 1. However, for our mobile EEG tasks (2 and 3), items did not appear and disappear but remained in the environment and could be seen as soon as they appeared in the participants’ field of view. As a consequence, items could be seen from a distance and approached, and perceived for extended periods or only fleetingly. This means we did not have an ‘event onset’ as such, and the concept of an evoked response did not clearly apply. Arguably, this more closely mirrors natural perception and behaviour, but presents challenges for data analysis. To overcome this issue, participants pushed a button when they arrived at, and were looking at the stimuli. Using this marker for the epoch-based analysis, we extracted oscillatory power for each trial over a 2 second period to test for power changes between our conditions without requiring a time-locked point. However, we will also likely miss numerous times when each item was fixated upon, and when the item was first seen. An alternative, more flexible approach, could involve the use of eye-tracking to define the time-point when a stimulus is fixated upon, with this information being used to construct fixation-related potentials (Kamienkowski et al., 2012; Kristensen et al., 2017). As such, future studies could look to incorporate eye-tracking measures during mobile EEG and AR for more precise object recognition effects.

While this approach was successful for our epoch-based analysis, alternative approaches are needed to take advantage of the continuous nature of perception, where multiple different events occur distributed over time. In our GLM-based analysis, we demonstrate an approach to relate time-varying neural signals to time-varying measures of perception, again based on face inversion. Using the continuous processed EEG signals, rather than discrete epochs, we were successful in relating the time-varying perceptual characteristics of the visual environment to the time-varying neural characteristics measured with EEG. We achieved this using an GLM to relate EEG signals to a continuous measure capturing face inversion, with additional regressors for motion parameters of no interest. With this analysis we replicated the face inversion findings of the epoch-based approach, where the effect sizes were comparable across the analyses as were the topographic distributions of effect sizes. Although this was a simplified example, the GLM approach appears effective for studying relationships between the visual world and neural signals, allowing greater flexibility to relate continuous perpetual events to continuous EEG signals. This approach followed similar applications in fMRI (e.g. Huth et al., 2012) and MEG (e.g. Brodbeck et al., 2018) but in this case expanded the application to mobile, freely moving settings.

The use of AR in mobile EEG experiments designed to study perception and cognition is a novel approach. Unlike VR, where both objects and the environment are computer generated, AR allows for virtual objects to be embedded into complex real-world environments, opening up new possibilities of controlled experimental designs in natural settings. For example, it allows for the navigation of personally familiar routes taken on a frequent basis whilst changing the items seen on the route or the path taken. One question that becomes relevant when considering AR (or VR), is whether virtual objects elicit neural responses similar to seeing real objects. A similar question has been raised concerning neuroimaging studies based on 2D images of objects compared to perceiving real 3D objects, with sometimes different effects across the conditions (Snow et al., 2011). In the current study we used 3D virtual faces that resemble human faces, but did not have all the characteristics of natural faces. Yet our results demonstrate similar neural responses to virtual faces and to images of faces, demonstrating that AR is a feasible option for presenting 3D stimuli that can be reliably studied with EEG.

Whilst our study could have been conducted without AR, we believe that the combination of AR and mEEG provides a powerful paradigm for psychology and cognitive neuroscience. With increasingly complex tasks and environments, it becomes more important to retain some experimental control to both provide participants with similar experiences and enable the researchers to modify perception in a flexible manner. The increasing use of AR for experiments could allow more research to be conducted outside the lab, where participants retain a sense of agency and immersion in the environment. Current research suggests AR is more engaging than VR, with potential improvements in memory encoding (Maidenbaum et al., 2019), and is sensitive to changes in attentional states (Vortmann et al., 2019). The expanded use of AR and EEG presents a new tool for psychologists to meld virtual and real environments in naturalistic paradigms.

In conclusion, here we show that cognitively relevant neural signals can be detected in AR and mobile EEG paradigms. Similar to lab-based effects, we showed inverted AR faces elicit greater low frequency power compared to upright AR faces while participants freely moved through an indoor office space. The combination of AR and mobile EEG could offer a new paradigm for cognitive neuroscience, whereby cognition can be studied while participants are immersed in natural environments yet experimenters can retain some control over what items people see and how.

## Acknowledgements

This work was supported by a Royal Society and Wellcome Trust Sir Henry Dale Fellowship (211200/Z/18/Z). For the purpose of open access, the author has applied a CC BY public copyright licence to any Author Accepted Manuscript version arising from this submission. The authors report no conflict of interest.

